# Opponent neurochemical and functional processing in NREM and REM sleep in visual learning

**DOI:** 10.1101/738666

**Authors:** Masako Tamaki, Zhiyan Wang, Tyler Barnes-Diana, Aaron V. Berard, Edward Walsh, Takeo Watanabe, Yuka Sasaki

## Abstract

Sleep is beneficial for learning. However, whether NREM or REM sleep facilitates learning, whether the learning facilitation results from plasticity increases or stabilization and whether the facilitation results from learning-specific processing are all controversial. Here, after training on a visual task we measured the excitatory and inhibitory neurochemical (E/I) balance, an index of plasticity in human visual areas, for the first time, while subjects slept. Off-line performance gains of presleep learning were associated with the E/I balance increase during NREM sleep, which also occurred without presleep training. In contrast, increased stabilization was associated with decreased E/I balance during REM sleep only after presleep training. These indicate that the above-mentioned issues are not matters of controversy but reflect opposite neurochemical processing for different roles in learning during different sleep stages: NREM sleep increases plasticity leading to performance gains independently of learning, while REM sleep decreases plasticity to stabilize learning in a learning-specific manner.

## Main Text

An accumulating body of evidence demonstrates that sleep is beneficial for various types of learning and memory. However, the underlying neural mechanisms of how sleep impacts learning are poorly understood. At present, there are at least three serious controversies. The first controversy concerns the roles of sleep stages in the facilitation of learning. Sleep is largely categorized into non-rapid eye movement sleep (NREM) sleep and rapid eye movement (REM) sleep^1^. Some research groups have suggested that NREM sleep^2–5^ plays an important role in facilitating learning, while other groups emphasized the role of REM sleep^6–10^. Whether NREM and/or REM sleep contributes to learning is unclear^11–14^. The second controversy concerns how sleep is beneficial to learning. At least two different types of learning benefits have been noted: off-line performance gains of learning^3–5, 9, 15^ and resilience to interference^16–19^. Off-line performance gains mean that the performance of learning acquired before sleep is enhanced after sleep without any additional training. The resilience to interference by sleep refers to a decreased amount of retrograde interference, or disruption, with learning acquired before sleep by learning or training on a new task after sleep. Retrograde interference with learning is regarded as a manifestation of the very plastic learning state that is vulnerable to “rewriting” by subsequent new learning^20^. Elimination of retrograde interference by sleep suggests that learning acquired before sleep is stabilized by sleep. It is unclear how both of these benefits are associated with plasticity states occurring during sleep or how each sleep stage contributes to plasticity states. The third controversy is whether the facilitation of learning occurs in a learning-specific or learning-independent manner. The former assumes that only synapses or networks that are specifically involved in the acquisition of learning and memory before sleep are changed during subsequent sleep, which leads to performance improvement after sleep^12, 21, 22^. The latter is based on the synaptic homeostasis hypothesis, which is a use-dependent process^11, 23, 24^. In this theory, during wakefulness, synapses that are used are overly strengthened irrespective of learning. During subsequent sleep, these overall synaptic changes are downscaled for synaptic homeostasis^11, 23, 24^, and only stronger synapses survive.

To resolve these controversial issues, we used humans as subjects to take advantage of the fact that humans have significantly longer and more clearly distinguished NREM and REM sleep episodes than animals^1, 25^. Although it was regarded as difficult to examine neurochemical processing related to plasticity during sleep with humans, for the first time we successfully made simultaneous measurement of magnetic resonance spectroscopy (MRS) and polysomnography in sleeping humans as well as sophisticated psychophysics for visual perceptual learning (VPL), defined as the improvement on a visual task performance after visual experience^4, 26–29^. Specifically, we measured the concentrations of glutamate (Glx), an excitatory neurotransmitter, and gamma-aminobutyric acid (GABA), an inhibitory neurotransmitter in human early visual areas, using MRS concurrently with polysomnography that determines sleep stages. We chose early visual areas, because VPL has been found to be associated with changes in early visual areas^28, 30–32^, and measured the excitation (E)/inhibition (I) balance as the concentration of Glx divided by the concentration of GABA. The E/I balance is regarded as a reliable index of the degree of plasticity and stability in early visual areas^30, 33, 34^. The visual critical period in rodents^33^ starts with a relatively high E/I balance in early visual areas and ends with a lower E/I balance. In human studies, a higher E/I balance is proportional to the degree of plasticity, as measured by psychophysics, and negatively correlated with the degree of stability of VPL in human early visual areas^30, 34^.

Measuring the E/I balance during sleep, we found opponent neurochemical processing during NREM and REM sleep that was highly associated with plasticity and stability of VPL in human brains. Off-line performance gains and stabilization occurred during different sleep stages with opposite directions of functional and neurochemical processing. First, plasticity of early visual areas increased during NREM sleep, as shown by an increased E/I balance, which well predicted off-line performance gains. Importantly, the E/I balance increased during NREM sleep irrespective of whether learning occurred before sleep, indicating that plasticity increases in early visual areas during NREM sleep occurred irrespective of whether learning/training had occurred before sleep. Second, REM sleep following NREM sleep plays a role in stabilization, making learning before sleep resilient to retrograde interference by new learning after sleep and protecting the performance gains developed during NREM sleep. Without REM sleep, the performance gains that would have occurred by NREM sleep were nullified by new learning after sleep. During REM sleep, the E/I balance, which increased during NREM sleep, decreased to a lower level than baseline measured during wakefulness in early visual areas. The decrease in E/I balance during REM sleep was correlated with the degree of resilience to retrograde interference in VPL. Importantly, the decrease in E/I balance during REM sleep was not observed without presleep learning and therefore is learning specific.

These results consistently suggest that in facilitating learning, NREM sleep and REM sleep play distinctive roles that are subserved by the opposite directions of functional and neurochemical processing. NREM sleep is involved in increased plasticity, whereas REM sleep is involved in stabilization, as shown by decreased plasticity. Increased plasticity during NREM sleep is not learning dependent, whereas the increased stabilization during REM sleep is learning dependent. Without REM sleep followed by NREM sleep, performance gains cannot be stabilized and are retrogradely interfered with by learning of a new task after sleep. Thus, each of the opponent concepts such as performance gains vs. stabilization, contribution of NREM vs. REM sleep, and learning independent vs. learning dependent may not be issues of controversy but properties in one of two distinctive functions.

## Results

In Experiment 1, we examined the roles of NREM and REM sleep in the two possible roles of sleep facilitation of performance gains and stabilization of learning. To accomplish this goal, we tested how the E/I balance in early visual areas, which is regarded as an index of plasticity, changes across NREM and REM sleep and how it is related to performance gains and stabilization. In particular, we tested the opponent processing hypothesis that performance gains and stabilization of learning are subserved by opponent neurochemical processing during NREM and REM sleep, respectively. This hypothesis was inspired by the results of studies of VPL during wakefulness^30, 34–36^. These studies examined the mechanisms of performance improvement as a result of training and stabilization that prevents retrograde interference by new learning during wakefulness. In these studies, retrograde interference was found during wakefulness if the interval between two trainings was within a few hours. Such interference is attributed to the incomplete stabilization during wakefulness of the first learning.

Retrograde interference is no longer observed until after a few hours of wakefulness. Many researchers believe that this result is observed because soon after training for the first task, the state of the learning is still highly plastic, and the passage of a few hours is necessary to stabilize learning^20, 30, 34, 36, 37^. In that case, the brain should enter a plasticity state during and right after training and then become less plastic as stabilization proceeds. The E/I balance increased after training and decreased as stabilization proceeded to the baseline^30, 34^. These results of learning states <u>during wakefulness</u> in previous studies support the current hypothesis: if plasticity increases during sleep for learning acquired before sleep, this should be followed by stabilization, which is associated with a decrease in enhanced plasticity for learning to survive retrograde interference by new learning after sleep upon waking. To actualize such plasticity increases followed by stabilization during sleep, NREM sleep and REM sleep that constantly follows NREM sleep may be suitable. If this hypothesis is correct, the following opponent processing should be revealed during NREM and REM sleep: during NREM sleep after training on a visual task, the E/I balance in early visual areas should increase in correlation with off-line performance gains that are observed after sleep, whereas during REM sleep, the E/I balance should decrease and the resilience of retrograde interference with learning acquired before sleep by new learning after sleep should be negatively correlated with the E/I balance during REM sleep. Moreover, if REM sleep is necessary for stabilization for off-line performance gains, off-line performance gains should be observed only with subjects who had both NREM and REM sleep in the face of retrograde interference.

To test the above hypothesis, we trained 10 subjects (see **Supplementary Table 1** for the mean age) on two different texture discrimination tasks (Task A and Task B) before and after sleep (see **Fig. 1a** see **Methods** for more details). There were four test sessions (**Fig. 1b**): before training on Task A (pretraining test), after training and before sleep (posttraining A test), after sleep and before training on Task B (postsleep test), and after training on Task B (posttraining B test). In each test session, the performance of Task A was measured. The details of the task properties and the rationale of the experimental design are explained as follows.

**Fig. 1.**
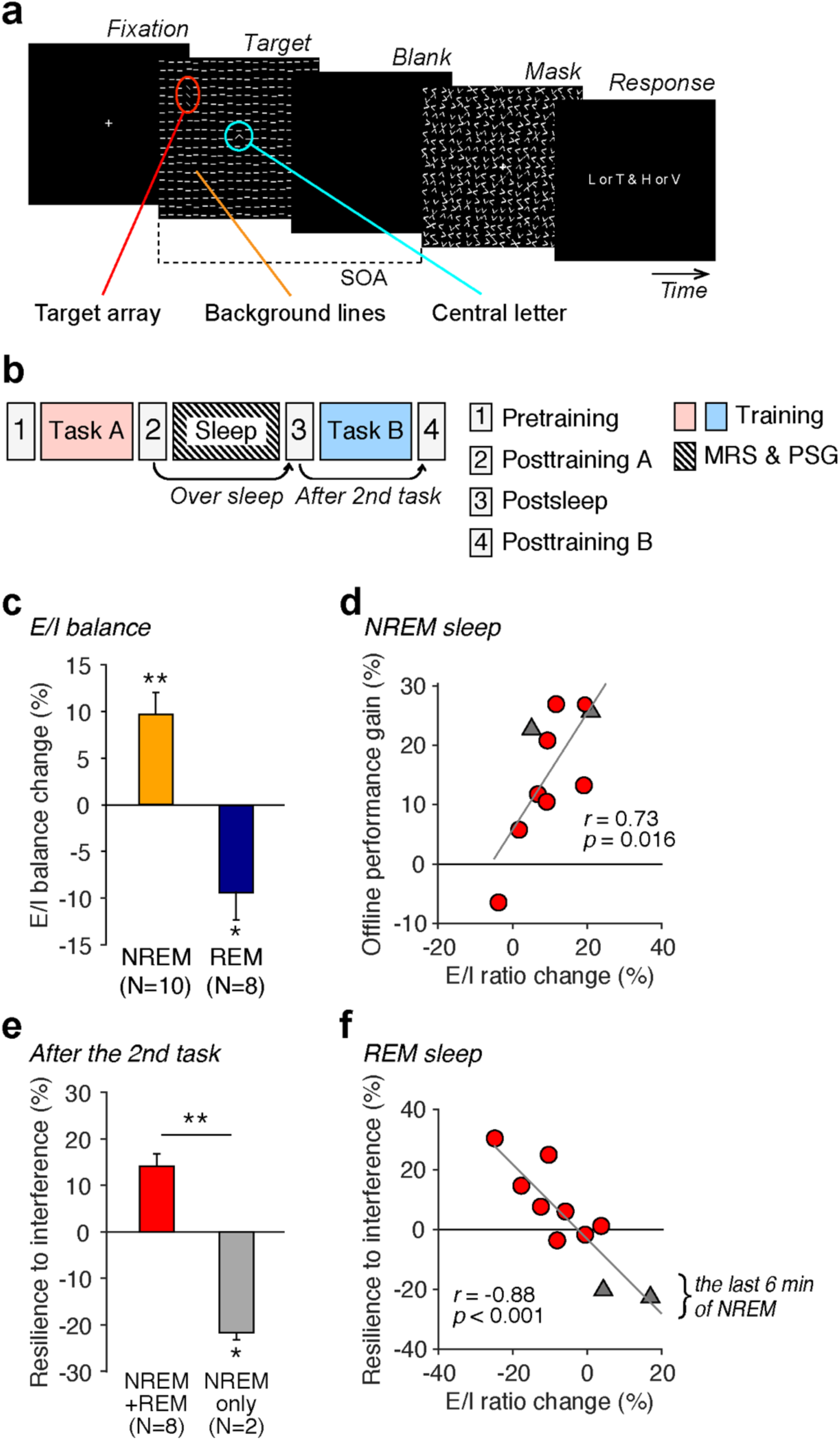
Procedures and results of Experiment 1. **a**, A TDT trial. One trial consisted of 5 displays: a fixation display, a target display, a blank display, a mask display and a response display. In the target display, there were three components: the background, target array and central letter. The orientation of the background line segments was either horizontal or vertical. A target array, which contained 3 line segments whose orientation was tiled from 45 deg from the orientation of the background segment, was presented in either the upper left or upper right visual field quadrant. The visual quadrant where the target array was presented was consistent throughout the experiment for each subject. The central letter was either ‘L’ or ‘T’. The purpose of the letter task was to ensure the subjects’ eyes fixation. After the mask, subjects were asked to respond whether the central letter of the target display was ‘L’ or ‘T’ and whether the orientation of the target array was ‘vertical’ or ‘horizontal’. A stimulus-to-mask onset asynchrony (SOA) was varied from trial to trial. See **Methods** for more details. **b**, Experimental design. Subjects were trained on the first task (Task A) before a nap and the second task (Task B) after the nap. **c**, The mean (± SEM) E/I balance changes from baselines to NREM sleep (n = 10) and REM sleep (n = 8). **d**, Scatter plots of E/I balance changes from baseline to NREM sleep vs. the off-line performance gains (the performance changes from the posttraining A to postsleep test sessions on Task A) for the NREM+REM group (red circles) and NREM only group (gray triangles). **e**, The mean (± SEM) degree of resilience to retrograde interference (performance change on first task from the postsleep to the posttraining B test sessions) for the NREM+REM and NREM only groups. **f**, Scatter plots of the E/I balance during REM sleep vs. the mean (± SEM) degree of resilience to retrograde interference for the NREM+REM group (red circles) and NREM only group (gray triangles). For the NREM only group, the E/I balance was obtained from the last 6 min of NREM sleep, as the average duration of REM sleep in the NREM+REM group was approximately 6 min. The correlation coefficient was *r* = -0.88 with all 10 subjects (*p* < 0.001), and *r* = -0.77 with only the NREM+REM group (n = 8, *p* = 0.025).

A TDT is a standard task for VPL^38^ and for research on the role of sleep in learning^3, 4, 9, 14, 39^. The TDT stimulus consisted of a target array of three-line segments aligned vertically or horizontally along with the background line segments. Subjects were repeatedly asked to report whether the orientation of the target array was horizontal or vertical. The performance was the threshold stimulus-to-mask onset asynchrony (SOA) of a task at which subjects show 80% correct responses. The orientations of the background elements in Task A and Task B were orthogonal to each other in the current experiment.

Learning TDT occurs specifically in the visual field quadrant where the target array is embedded in the background orientation^38, 39^. If two trainings on TDT with orthogonal background orientations as those in Task A and Task B are conducted sequentially, even though the targets are presented in the same visual field quadrant, the second training disrupted or interfered with learning the first training^35^. Thus, in the current experiment, we conducted training with Task A followed by Task B with sleep between the tasks to examine whether off-line performance gains were stabilized during sleep to test the feasibility of the opponent processing hypothesis.

During the sleep session (**Fig. 1b**), subjects slept inside the MRI scanner while polysomnography was performed (See **Methods** for more details and **Supplementary Table 2** for sleep structures) to objectively determine sleep stages. During NREM and REM sleep, Glx and GABA concentrations were measured from early visual areas (see **Supplementary Fig. 1** for an example location) using MRS, and the average Glx and GABA concentrations were computed for each sleep stage (see **Methods**). See **Supplementary Fig. 2** for example spectra from early visual areas.

First, to examine the amount of off-line performance gains on the first task from before to after sleep, we measured the performance change between the posttraining A test session before sleep and postsleep test session after sleep (**Fig. 1b**). The degree of performance gains was defined by [(threshold SOA in posttraining A test – threshold SOA postsleep test) / threshold SOA in posttraining A test] x 100%, where threshold SOAs were for Task A. The degree of performance gains was significantly greater than zero (one sample *t*-test, *t* (9) = 2.95, *p* = 0.016, Cohen’s *d* = 0.93).

Then, we examined the relationship between the degree of off-line performance gains and E/I balances during NREM and REM sleep relative to baselines that were measured during wakefulness (for the sake of simplicity, we will call them *E/I balances*) in early visual areas. First, as in **Fig. 1c**, the E/I balance increased during NREM sleep (one sample *t*-test, *t* (9)=3.94, *p* = 0.003, *d* = 1.25), while it decreased during REM sleep (one sample *t*-test, *t* (7) = 2.90, *p* = 0.023, *d* = 1.02) compared to that at baseline. Second, performance gains were significantly correlated with the E/I balance during NREM sleep (*r* = 0.82, *p* < 0.001), as shown in **Fig. 1d**, whereas it was not significantly correlated with the E/I balance during REM sleep (*r* = 0.60, *p* = 0.116). These results are in agreement with one of the hypothesized opponent processes in which off-line performance gains are related to the E/I balance increase, particularly during NREM sleep.

Then, we examined the second part of the opponent processes hypothesis. That is, we examined how retrograde interference is, if at all, reduced due to stabilization during sleep between the two trainings. As shown in **Fig. 1b**, if performance on Task A tested after the training on Task B was worse than after sleep and before the training on the 2nd task, this should indicate the retrograde interference from the training on Task B on learning of Task A. Here, we defined the degree of performance change on Task A from before and after the training on Task B by [(threshold SOA in postsleep test – threshold SOA in posttraining B test / threshold SOA in postsleep test] x 100%, where the threshold SOAs are for Task A. If the degree of performance change is lower than zero, this indicates retrograde interference. Thus, the degree of performance change from the postsleep test session to the posttraining B test session is proportional to the degree of resilience to the retrograde interference.

As stated above, to address the question of whether REM sleep is necessary for stabilization for off-line performance gains, we analyzed the results by dividing subjects into two groups (**Fig. 1e**). Eight out of ten subjects showed both NREM and REM sleep (NREM+REM group), while the remaining two subjects showed only NREM sleep without REM sleep (NREM only group). First, **Fig. 1e** shows that the degree of resilience to retrograde interference is significantly larger for the NREM+REM group than for the NREM only group (unpaired t-test, *t* (8) = 2.56, *p* = 0.009, *d* = 3.52). Second, **Fig. 1f** shows that the E/I balance (the x-axis) was below zero in 7 out of 8 subjects from the NREM+REM group and was also significantly negatively correlated with the degree of resilience across the 8 subjects (*r* = -0.77, *p* = 0.025, n = 8). Third, the performance of Task A for the NREM only group also increased during NREM sleep (gray triangles in **Fig. 1d**). However, Task A performance for the NREM only group significantly decreased after training on Task B (gray triangles in **Fig. 1f**). This decrease contributed to the extremely high correlation between the degree of resilience to the retrograde interference and the E/I balance measured during the last part of the sleep period (the E/I balance during REM sleep for the NREM+REM group and the E/I balance during the last part of NREM sleep for the NREM only group) with all subjects (**Fig. 1f**, *r* = -0.88, *p* < 0.001, n = 10; if we consider only the NREM+REM group, *r* = - 0.77, *p* = 0.025, n = 8). Fourth, the results of control measurements and analyses indicate that sleepiness (**Supplementary Table 3**) and the initial performance levels (**Supplementary Table 4**) were not significantly different between the NREM+REM group and the NREM only group. Thus, the difference in the resilience to retrograde interference between these groups are unlikely to be due to the difference in sleepiness or the level of initial performance. These results consistently suggest two distinctive mechanisms as follows: during NREM sleep, plasticity becomes greater, which leads to off-line performance gains after sleep. During REM sleep, plasticity becomes lower than that during wakefulness, which eliminates or reduces retrograde interference.

However, in Experiment 1, the number of subjects who had only NREM sleep was too small (n = 2). Thus, we conducted a behavioral experiment (Experiment 2) with a significantly larger number of subjects (n = 30; **Supplementary Table 1** for the mean age). As in Experiment 1, two trainings on different tasks were conducted with sleep between them. Performance gains were measured by conducting the pretest before the first training and the posttest after the second training (**Fig. 2a**). Polysomnography was applied to determine sleep stages.

**Fig. 2.**
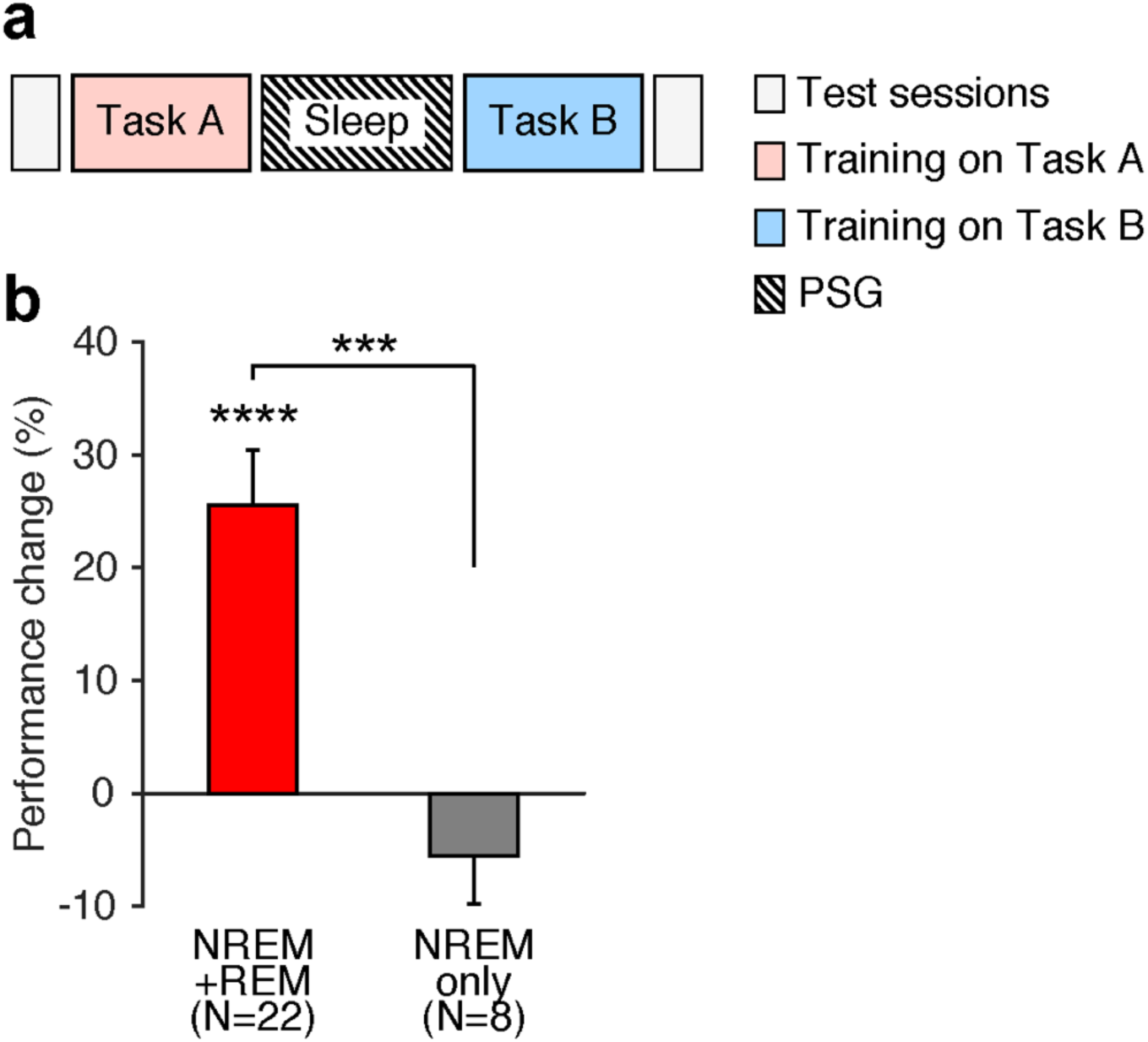
The experimental design and results of Experiment 2. **a**, Design. The first TDT training (Task A) was conducted with background A, and the second training (Task B) was conducted with background B. Test sessions were conducted before the first training (pretest) and after the second training (posttest) to measure performance gains on the first training (Task A). **b**, Mean (± SEM) performance change (± SEM) for the first training (Task A) from the pretest session to the posttest session in the REM-present (red, n = 22) and REM-absent (gray, n = 8). \*\*\*\**p* < 0.001, \*\*\**p* < 0.005.

Twenty-two subjects had both NREM sleep and REM sleep, whereas the remaining 8 subjects had only NREM sleep among 30 subjects who slept between the two trainings. As shown in **Fig. 2b**, significant off-line performance gains were observed among subjects who had both NREM sleep and REM sleep (one sample *t*-test, *t* (21) = 5.24, *p* < 0.001, *d* = 1.12), whereas the subjects who had only NREM sleep did not show any significant off-line performance gains (one sample t-test, *t* (7) = 1.30, *p* = 0.236). Therefore, the off-line performance gains were significantly different between the groups (unpaired *t*-test, *t* (28) = 3.63, *p* = 0.001, *d* = 1.70). These results are consistent with those in Experiment 1 and indicate that the increased plasticity state during NREM sleep needs to be stabilized during REM sleep to retain off-line performance gains and allow learning to survive retrograde interference.

Then, a question arises: do plasticity changes in early visual areas during NREM and REM sleep occur only after learning/training or are the plasticity changes innate with sleep itself? In other words, are the E/I balance increase during NREM sleep and/or the E/I balance decrease during REM sleep as shown in Experiment 1 learning specific or not? If a change is learning specific, the E/I balance should increase during NREM sleep and/or it should decrease during REM sleep only after training on a visual task. If a plasticity change is not learning dependent, the E/I balance increases during NREM sleep, and/or its decreases during REM sleep should occur irrespective of whether learning/training occurred before sleep. To address this question, subjects (n = 9, see **Supplementary Table 1**) participated in Experiment 3 with the same procedures as those in Experiment 1, except that no training was conducted before or after sleep. The circadian timings of the nap in Experiments 1 and 3 were equivalent (see **Methods**).

**Fig. 3a** shows that both results of Experiment 1 in which learning occurred and those in Experiment 3 in which no learning occurred. First, in Experiment 3, six subjects showed NREM sleep followed by REM sleep (NREM+REM group), while the remaining three subjects did not (NREM only group). The mean E/I balance in early visual areas was significantly higher during NREM sleep than at baseline (one sample *t*-test, *t* (8) = 4.71, *p* = 0.002, *d* = 1.57) in all subjects in Experiment 3. This result is consistent with the results of Experiment 1 in which learning occurred. In contrast, during REM sleep, the E/I balance was not significantly different from the baseline (one sample *t*-test, *t* (5) = 2.12, *p* = 0.087). This is different from the results of Experiment 1, which showed that the E/I balance was significantly lower than zero during REM sleep (one sample *t*-test, *t* (7) = 2.90, *p* = 0.023, *d* = 1.02, as shown in Fig. 1c).

**Fig. 3.**
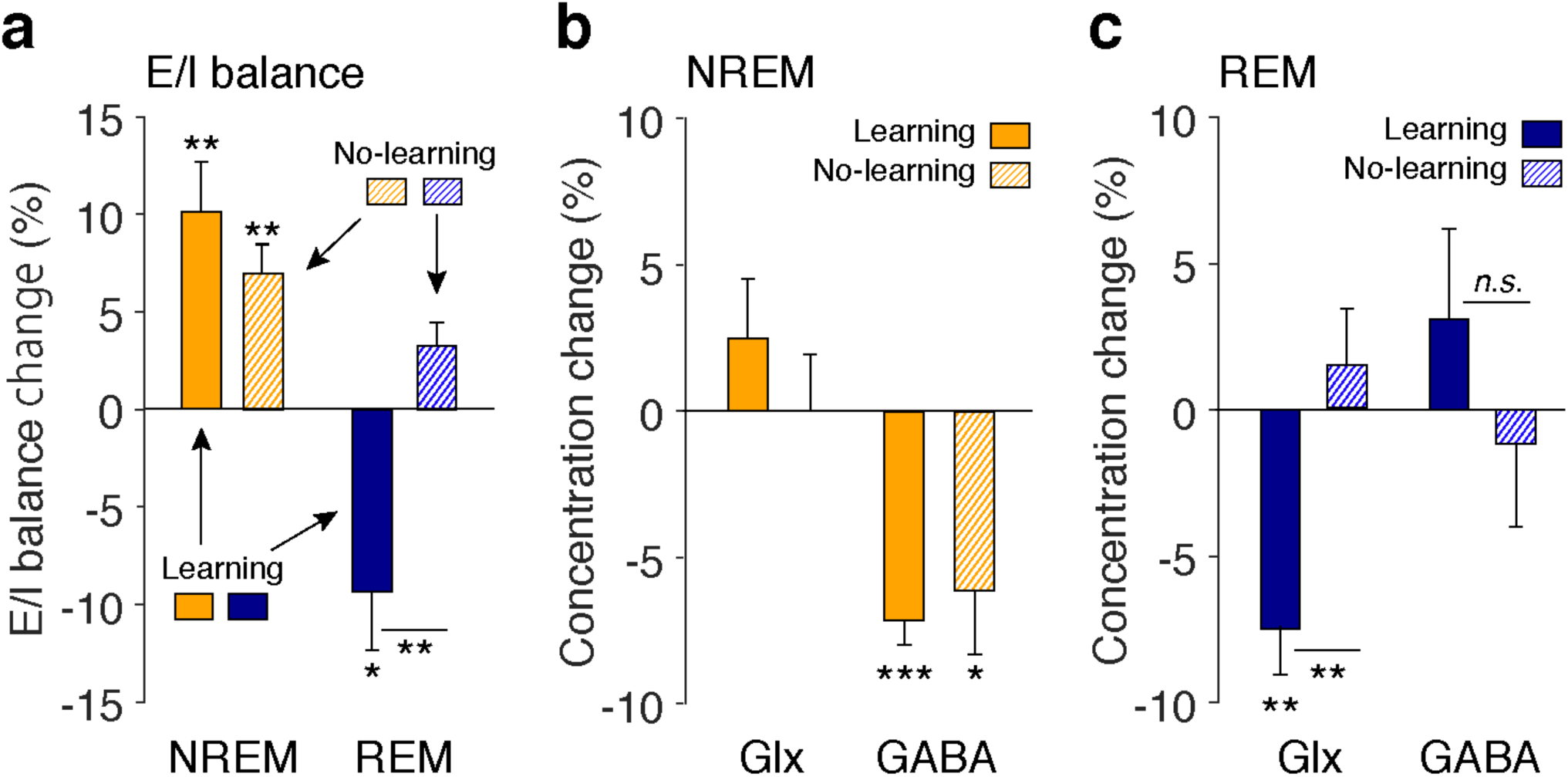
Collective results of Experiments 1 and 3. **a**, E/I balance during NREM and REM sleep after learning (n = 10 for NREM sleep, n = 8 for REM sleep, the same as **Fig. 1c**) in Experiment 1 and after no prior learning (the control condition, n = 9) in Experiment 3. **b**, The concentrations of Glx and GABA during NREM sleep. **c**, The concentrations of Glx and GABA during REM sleep. \*\*\**p* < 0.005, \*\**p* < 0.01, \**p* < 0.05.

Second, we directly compared the results of Experiments 1 and 3 by two-way ANOVA with factors of Learning (learning presence vs absence) and Sleep Stage (NREM vs. REM sleep) on the E/I balance. The results showed a significant interaction of Learning x Sleep Stage factors (*F* (1, 28) = 10.46, *p* = 0.003) as well as a significant main effect of Sleep Stage (*F* (1, 28) = 22.29, *p* < 0.001). A post hoc analysis indicated that the E/I balance during REM sleep was significantly different between learning presence (Exp. 1) vs. absence (Exp. 3) conditions (unpaired *t*-test, *t* (12) = 3.16, *p* = 0.008, *d* = 1.80), whereas no significant difference in the E/I balance was found during NREM sleep between these conditions (unpaired *t*-test, *t* (17) = 1.02, *p* = 0.320). Finally, the E/I balance was significantly lower than zero during REM sleep after learning occurred (Exp. 1, one sample t-test, *t* (7) = 2.90, *p* = 0.002, *d* = 1.57), while it showed no significant difference from zero during REM sleep when learning did not occur (Exp. 3, one sample t-test, *t* (5) = 2.12, *p* = 0.087). These results confirmed that the E/I balance of early visual areas during NREM sleep increased irrespective of whether learning occurred (Exp. 1) or did not occur (Exp. 3), while the E/I balance during REM sleep decreased relative to the baseline only when learning occurred (Exp. 1).

As discussed above, the E/I balance is determined by the concentration of Glx divided by the concentration of GABA in early visual areas. What roles do these neuromodulators play in off-line performance gains and plasticity enhancement during NREM sleep and resilience to retrograde interference and stabilization during REM sleep? As shown in **Fig. 3b**, in both Experiment 1 in which learning occurred and Experiment 3 in which no learning occurred, during NREM sleep, the concentration change in GABA was significantly lower than zero (one sample *t*-tests, Exp. 1, *t* (9) = 9.22, *p* < 0.001, *d* = 2.92; Exp.3, *t* (8) = 2.81, *p* = 0.023, *d* = 0.94), while the concentration change in Glx was not significantly different from zero with (one sample *t*-test, Exp. 1, *t* (9) = 1.23, *p* = 0.249) or without learning (Exp. 3, *t* (8) <0.01, *p* = 0.9994). Thus, the decreases in GABA concentration seem to be a major factor of the E/I balance elevation during NREM sleep. Alternatively, during REM sleep, as shown in **Fig. 3c**, Glx concentration was significantly lower than zero only after learning (Exp. 1, one sample *t*-test, *t* (7) = 4.23, *p* = 0.004, *d* = 1.50), showing a significant difference between Experiments 1 and 3 (unpaired *t*-test, *t* (12) = 3.10, *p* = 0.009, *d* = 1.65). The decreases in Glx concentration seem to mainly determine the E/I balance decrease during REM sleep after learning. See **Supplementary Figs** for quality controls for MRS data, including NAA linewidth (**Supplementary Fig. 3**), and the concentration of NAA that was used as a reference metabolite (**Supplementary Fig. 4**). The series of quality control data for MRS (**Supplementary Table 5**), such as the shim value (Hz), NAA linewidth (Hz), frequency drift (Hz), and Cramer-Rao lower bound (%SDs for Glx and GABA, respectively), indicate that no significant difference in the MRS data quality between Experiments 1 and 3.

Thus far, we have assumed that the resilience of retrograde interference is a manifestation of stabilization of learning the first task by REM sleep. However, an alternative interpretation to stabilization of the first learning may possibly account for why the first remains after the second learning. Some researchers have put forth the resource consumption hypothesis in which the first learning consumes all of the plasticity resources during the sleep period, which does not lead to the second learning^40^. If this was the case, stabilization would not need to be assumed because there would be no second learning that interferes with the first learning. Both the stabilization and resource consumption hypotheses predict the survival of the first learning over the training on the second task after sleep. However, a crucial difference in the predictions by these two possibilities lies in the second learning. The resource consumption hypothesis predicts that the second learning should not occur because a plasticity resource is consumed by then, while the stabilization hypothesis does not predict the lack of plasticity resources and thus predicts that the second learning should occur. To test which possibility is more plausible, we examined whether the first and second learning occurred after NREM and REM sleep and NREM sleep only in Experiment 4 (Fig. 4a). The procedures of this experiment were identical to those of Experiment 2, except that in the pretest and posttest, performances of the second task as well as the first task were measured.

**Fig. 4.**
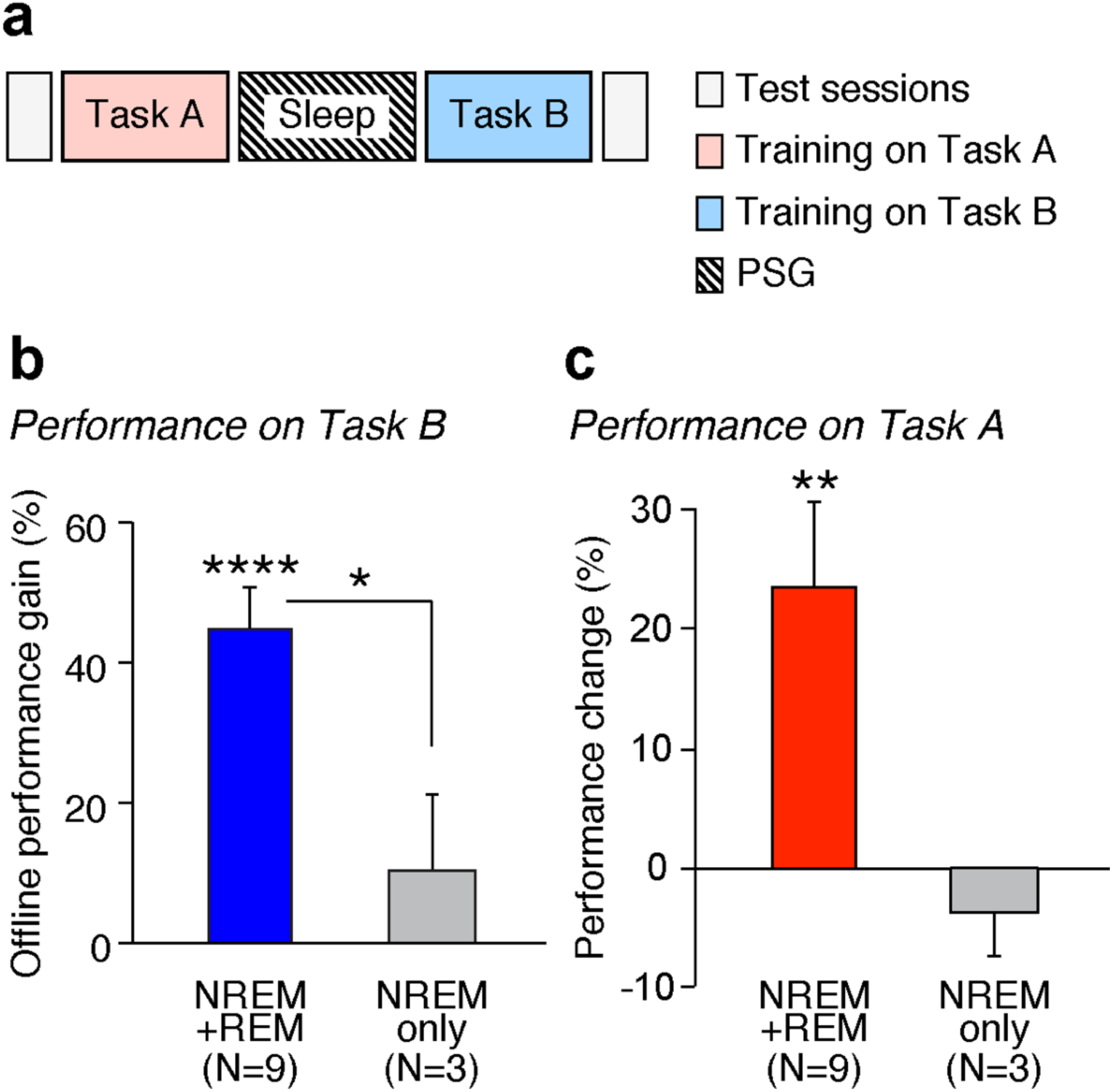
The experimental design and results of Experiment 4. **a**, Design. The first TDT training was conducted with background A (Task A, pink), and the second training was conducted with background B (Task B, cyan). Test sessions were conducted before the first training (pretest) and after the interval and the second training (posttest) to examine performance changes in both Tasks A and B. **b**, Mean performance change (± SEM) for Task B (background orientation B) at the posttest session from the pretest session for the group of subjects who had NREM and REM sleep (NREM+REM group, blue, n = 9) and the group who showed only NREM sleep (NREM only group, gray, n = 3). The performance change was significantly different from zero for the NREM+REM sleep group (\*\*\*\**p* < 0.001) and across the two groups (\**p* <0.05). **c**, Mean performance change (± SEM) for Task A at the posttest session from the pretest session with the same two groups as in B (one sample *t*-test against 0, NREM+REM, *t* (8) = 3.40, *p* = 0.009; NREM only, *t* (2) = 1.03, *p* = 0.411).

The results are consistent with the stabilization hypothesis. Similar to Experiment 1, the subjects were divided into the NREM+REM group (n = 9) and NREM only group (n = 3), depending on whether REM sleep appeared during the interval. The stabilization hypothesis predicts that the NREM+REM group learns Task B together with stabilization of Task A, while the NREM only group does not learn Tasks A and B due to lack of stabilization during REM sleep. In contrast, the plasticity resource hypothesis predicts that both groups learn Task A but not task B. We tested whether the performance change was significantly different by a 2-way ANOVA with Group (NREM+REM vs. NREM only) and Task (first A vs. second B) factors. The results show significant main effects of Group (*F* (1, 10) = 8.258, *p* = 0.017) and Task (*F* (1, 10) = 8.518, *p* = 0.015) without an interaction (*F* (1, 10) = 0.395, *p* = 0.544). The results indicate that the performance of both tasks was significantly different between the groups. The NREM+REM group had significant performance gains for Task B (one sample *t*-test, *t* (8) = 7.51, *p* < 0.001, *d* = 2.50, the blue bar in **Fig. 4b**) as well as for Task A (one sample *t*-test, *t* (8) = 3.40, *p* = 0.009, *d* = 1.13, the red bar in **Fig. 4c**). In contrast, the NREM only group had no performance gains for Task A or Task B (one sample *t*-test, Task A, *t* (2) = 1.03, *p* = 0.411; Task B, *t* (2) = 0.95, *p* = 0.441, gray bars in **Fig. 4b, c**).

The occurrence of the second learning in the NREM+REM group is not in agreement with the consumption hypothesis, while it is in agreement with the stabilization hypothesis. The absence of the first and second learning in the NREM only group also contradicts the consumption hypothesis because the consumption hypothesis predicts that the second learning should occur if the first learning does not occur and thus does not use the plasticity resource. These results consistently support that stabilization occurs during the NREM+REM group, while learning remains fragile in the NREM only group without REM sleep but not in accordance with the resource consumption hypothesis.

## Discussion

In this study, we examined the opponent process hypothesis, that is, roles of NREM and REM sleep in VPL by measuring the concentrations of neurotransmitters such as Glx and GABA in human early visual areas as well as polysomnography and behavioral measurements. This study was the first to measure neurotransmitters in human subjects during sleep using MRS. We calculated the E/I balance since it is a good index of the degree of visual plasticity in humans and animals^30, 33, 34^. Consequently, we made several important findings. First, performance on the task learned before sleep was increased and resilient to the subsequent training on a new task if the sleep contained both NREM and REM sleep. The amount of such off-line performance gains after sleep was highly correlated with the E/I balance during NREM sleep relative to the baseline measured after training during wakefulness. However, the amount of off-line performance gains was not significantly correlated with the E/I balance during REM sleep relative to baselines. Second, off-line performance gains processed and obtained for NREM sleep did not stabilize without REM sleep. Third, the E/I balance significantly increased during NREM sleep after no training as well as after training. Fourth, retrograde interference was not observed following new training after both NREM and REM sleep, whereas retrograde interference occurred following new training only after NREM sleep. Fifth, the degree of resilience to retrograde interference was inversely correlated with the E/I balance during REM sleep, while it was not significantly correlated with the E/I balance during NREM sleep. Finally, the reduction in retrograde interference after sleep consisting of NREM and REM sleep is likely due to stabilization during sleep rather than plasticity resource consumption.

These results suggest two distinctive mechanisms for NREM sleep and REM sleep with opponent patterns of neurochemical processing, as shown in **Fig. 5 and Table 1**. One mechanism is plasticity increases in early visual areas during NREM sleep. The plasticity increases with E/I balance enhancement, irrespective of whether learning occurred before sleep. The other mechanism is stabilization. During REM sleep, learning is stabilized by decreasing once increased plasticity during NREM sleep, as shown by decreases in the E/I balance below baseline. In contrast to plasticity increases during NREM sleep, the decreases in the E/I balance below baseline occur only after learning occurred during REM sleep, that is, in a learning-dependent manner. If stabilization-induced E/I balance decreases does not occur after plasticity increases during NREM sleep (not followed by REM sleep), no off-line performance gains occur after the second learning. This finding indicates that NREM sleep and REM sleep coordinate to induce and stabilize off-line performance gains.

**Fig. 5.**
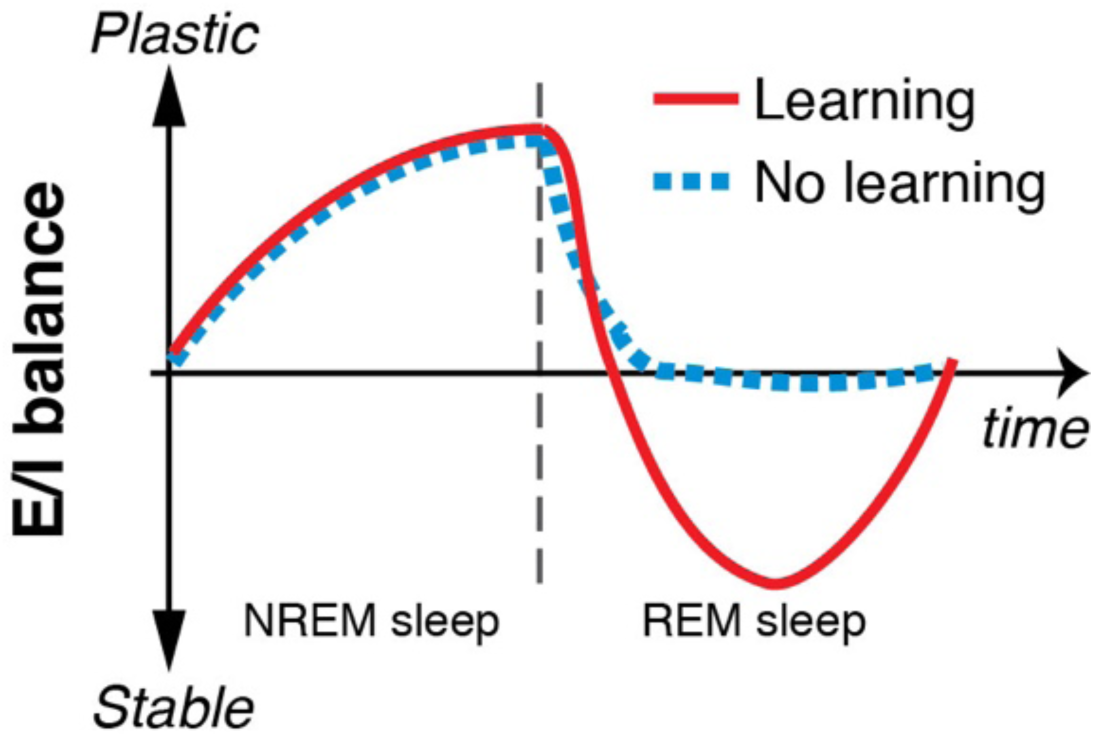
An illustration of the opponent process hypothesis in terms of plasticity/stability modulation (E/I balance) during NREM and REM sleep with or without presleep learning as a function of time.

**Table 1.**
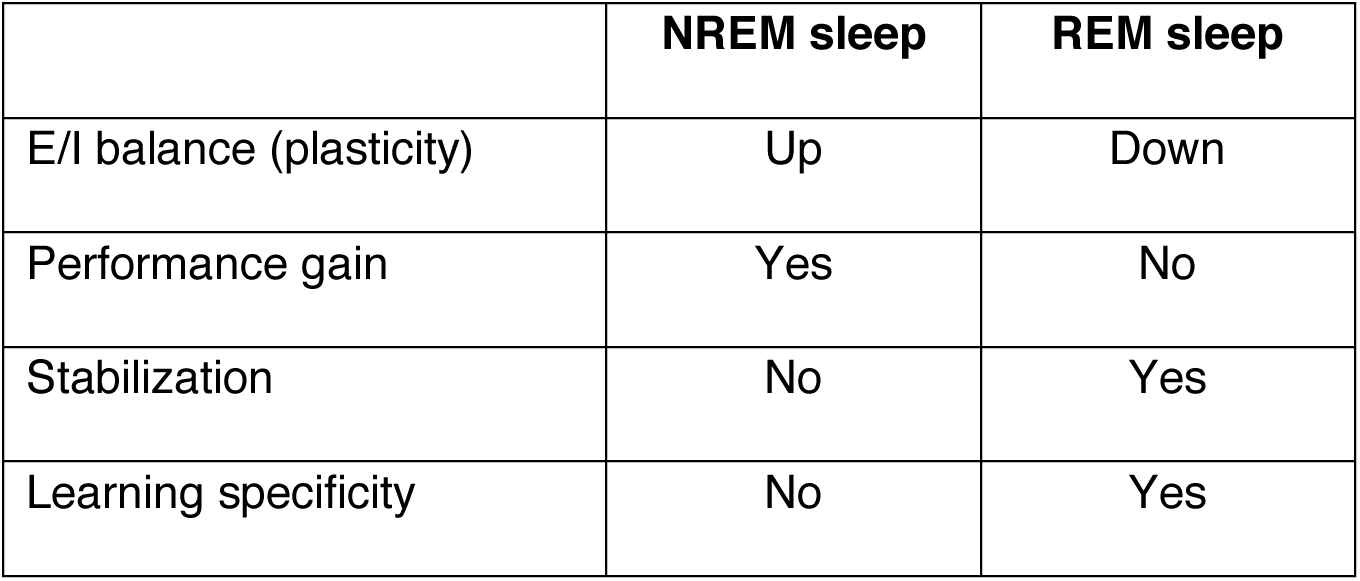
Functional and neurochemical differences between NREM and REM sleep

|  | **NREM sleep** | **REM sleep** |
| --- | --- | --- |
| E/I balance (plasticity) | Up | Down |
| Performance gain | Yes | No |
| Stabilization | No | Yes |
| Learning specificity | No | Yes |

This opponent processing model may help resolve the abovementioned three controversies. Instead of assuming only one of the two roles, such as facilitating off-line performance gains and stabilization, sleep plays both roles in a complementary fashion. Instead of assuming that NREM or REM sleep should work for learning facilitation, NREM sleep likely plays a role in off-line performance gains, while REM sleep stabilizes plasticity increases during NREM sleep. Instead of assuming that sleep facilitation is exclusively learning dependent or not, a plasticity increase during NREM sleep occurs in a learning-independent manner, whereas stabilization during REM sleep may occur in a learning-specific manner.

The stabilization that occurs during REM sleep is distinguished from the typical stabilization during wakefulness in several manners. The typical stabilization process during wakefulness is associated with gradual decreases of once increased E/I balance due to training to the baseline E/I balance before training^30^. On the other hand, the stabilization during REM sleep is associated with the E/I balance that became even <u>lower than</u> baseline in a short time. The E/I balance lower than baseline was observed during wakefulness due to training that was continued even after performance increases reached an asymptote, called overlearning^30^. Overlearning is governed by hyperstabilization, which more strongly and rapidly stabilizes a learning state^30^. Thus, the stabilization during REM sleep after learning is not the same as the typical stabilization in which the E/I balance decreases to baseline in a few hours after learning but shares some aspects with hyperstabilization during wakefulness, which plays a role in rapidly and strongly stabilizing a learning state^30^.

E/I balance changes based on Glx and GABA measured in our experiments may reflect, if not entirely, synaptic transmission. We measured the concentrations of Glx and GABA in early visual areas using MRS. While MRS measures various types of metabolites outside and inside neurons, we argue that changes in Glx, which largely reflects glutamate^41^, and GABA within the same subjects caused by sleep stages and learning likely largely represent synaptic transmission of Glx and GABA for the following reasons. First, various manipulations including drug intakes^42^, visual stimulation^43^, transcranial direct current or magnetic stimulation^44–49^ and memory tasks^41, 50, 51^ modulate glutamatergic and GABAergic synaptic transmissions, which are shown as changes in the Glx and GABA concentrations measured by MRS. As these manipulations may modulate synaptic transmission, the changes in Glx and GABA measured in vivo by MRS in the present study may also largely reflect synaptic transmission. Second, we measured Glx and GABA concentrations in early visual areas. The change in the concentrations of these neurotransmitters in early visual areas measured by MRS in the current study was consistent with that in previous studies^30, 34^, which was highly correlated with the degree of plasticity or stability of VPL^30, 34^. Third, in our experiments, the Glx and GABA concentrations in early visual areas were compared between within-subject conditions that differed only in learning and/or sleep stages, with other factors held constant. Changes in Glx and GABA in different learning and/or sleep conditions largely came from synapses^52–58^. Thus, we suggest that concentration changes found in the present study may, if not entirely, reflect synaptic transmission changes.

The decreases in GABA concentrations in early visual areas during NREM sleep may seem counterintuitive, given that subcortical GABAergic neurons inhibit the ascending arousal system and initiate sleep^59, 60^. One possible explanation is that the activity of these subcortical GABAergic neurons not only leads to sleep but also inhibits local inhibitory interneurons in early visual areas^33, 53, 61^. These disinhibited neurons could enhance plasticity in early visual areas, which leads to off-line performance gains when visual learning occurs before sleep.

The learning-dependent decrease in Glx during REM sleep may be associated with reduction in glutamatergic excitatory synapses in activated neurons during preceding NREM sleep in early visual areas. Although it is beyond our scope to speculate the underlying detailed molecular mechanisms, our speculation is based on the following reasons. REM sleep selectively prunes and maintains new synapses in learning^10^. Such models of pruning and maintenance of synapses in early visual areas are consistent with the role of REM sleep as stabilization of the first visual learning and permission of the second visual learning were found in the current study. We assume that the synapses that are involved in visual learning are reactivated during NREM sleep^62^, and most important synapses of them will be maintained and less important synapses will be pruned during REM sleep. If these pruned synapses are glutamatergic, as excitatory synapses are considered to be a molecular basis of learning^58, 63, 64^, pruning of glutamatergic synapses may result in a reduction in the Glx concentration in early visual areas.

One may wonder whether a model of synaptic changes during sleep suggested by a recent review paper^61^ is directly related to our findings. Based on results of animal sleep studies, their model assumes opponent synaptic changes during sleep; local synaptic enhancement during slow wave sleep that occurs specifically for learning and global synaptic downscaling during REM sleep that occurs in a sleep-independent manner. Although this model is highly interesting, it concerns neural processing at a different level/dimension from the opponent functions we found and are not directly linked to our findings. First, local synaptic enhancement and global downscaling in this model do not have one-to-one correspondences to performance gain and stabilization in our findings, respectively. Note that the stabilization in our study is defined as the processing of making presleep learning resilient to retrograde interference by new postsleep learning^20^. It is unclear how resilience to interference is formed by their model. Second, in their model, local synaptic enhancement is assumed to occur in a learning-dependent manner, while global downscaling does not occur specifically for learning. On the other hand, we found that performance gain occurs in enhanced plasticity that occurs in learning-independent manner, while stabilization occurs in a learning-dependent manner. The model of opponent synaptic changes^61^ and our findings of performance gain vs stabilization may be on different dimensions/levels. A future study would clarify how performance gains and stabilization in our findings are related to local synaptic enhancement and global synaptic downscaling.

In summary, we examined the roles of NREM sleep and REM sleep in off-line performance gains and stabilization of VPL. To take advantage of measuring brain states of sleep in humans, which have significantly longer and clearer sleep cycles than most other species, we measured Glx and GABA in the sleeping human brain to obtain the E/I balance, which is an in vivo plasticity/stability index. The results are consistent with the opponent processing model in which off-line performance gains occur due to plasticity increases in early visual areas in a learning-independent manner during NREM sleep, while REM sleep plays a role in stabilizing the increased plasticity during NREM sleep in a learning-dependent manner and enhancing resilience to retrograde interference by new learning. Such opponent processing may lead to resolving the outstanding controversies surrounding the benefits of facilitation of learning, benefits of sleep for learning, involvement of sleep stages, and specificity of mechanisms for the facilitation of learning. This information will also lead to research that will examine how abnormalities in sleep patterns^65–71^ influence the proposed complementary roles of sleep in VPL and other types of learning and memory.

## Methods

### Participants

A total of 51 young healthy subjects participated in the present study. See **Supplementary Table 1** for the mean age and gender information. All subjects gave written informed consent for their participation in experiments. This study was approved by the institutional review board at Brown University. We removed 1 subject’s data from Experiment 2 and another subject’s data from Experiment 3 since these subjects showed sleep-onset REM sleep shortly after lights off. Thus, the total number of subjects analyzed was 49.

Subjects were required to have normal or corrected to normal vision. No subjects had medical conditions, including sleep disorders. All of the subjects had no prior experience in visual perceptual learning tasks. People who play action video games frequently were excluded because extensive video game playing affects visual and attention processing^72, 73^. In addition, participants were required to have a regular sleep schedule. Anyone who had a physical or psychiatric disease, was currently under medication, or was suspected to have a sleep disorder was not eligible to participate^74, 75^.

### Experimental designs

We first describe common procedures to all experiments and then procedures specific to each experiment.

#### Common procedures

Before the experiments started, subjects were instructed to maintain their regular sleep-wake habits, i.e., their daily wake/sleep time and sleep duration until the study was over. The sleep-wake habits of subjects were monitored by a sleep log for about a week prior to the experiment. In addition, all the subjects in Experiments 1 and 8 subjects in Experiment 3 were asked to wear a wrist actigraphy device (GT9X-BT, ActiGraph, LLC).

On the day before the experiment, subjects were instructed to refrain from alcohol consumption, unusually excessive physical exercise, and naps. For all experiments, caffeine consumption was not allowed on the day of experiments.

The sleep session started in the early afternoon (1 pm-2 pm) for all experiments.

In Experiments 1, 2, and 4, there were two training sessions for the texture discrimination task (TDT) (see ***Texture discrimination task*** and see ***SOAs and the number of TDT trials*** for the number of trials), which were separated by a 120-min interval. The background orientations of the first (Task A) and second (Task B) TDT tasks were orthogonal to each other (horizontal or vertical). During the 120-min interval, subjects had a 90-min sleep period and a 30-min break. The 30-min break was to ameliorate sleep inertia^76^. During the 30-min break, a questionnaire was administered to obtain subjects’ introspection about their sleep, including subjective sleep-onset latency, subjective wake time after sleep onset, comfort of the environment, and occurrence of dreams^75^.

In Experiments 1, 2 and 4, subjective (Stanford sleepiness scale; SSS)^77, 78^ and behavioral sleepiness (psychomotor vigilance test; PVT^79^) were measured prior to each test session for TDT (see ***Sleepiness measurement*** for more details). In Experiment 3, there was no test session for TDT as there was no learning involved in this experiment (see below). Four out of 10 subjects in Experiment 3 rated their sleepiness using SSS before electrode preparation, which was approximately 40 min before subjects entered the scanner (see ***Sleepiness Measurement***). This timing roughly matched when SSS was measured for the subjects in Experiment 1 in the posttraining A test session.

For all the experiments, there was an additional adaptation sleep session before the main sleep session. When subjects sleep in a sleep laboratory for the first time, the sleep quality is degraded due to the first-night effect (FNE) caused by the new environment^74, 75, 80–82^. The adaptation sleep session was necessary to mitigate the FNE. During the adaptation sleep session, all electrodes for PSG measurement were attached to the subjects (see ***PSG measurement*** below). Subjects slept in the same fashion as in the main sleep session, with (Experiments 1 and 3) or without (Experiments 2 and 4) MRI scans. The adaptation session was conducted approximately one week before the main experimental sleep session so that any effects due to sleeping during the adaptation nap would not carry over to the experimental sleep session.

#### Experimental design for Experiment 1

There were four test sessions: pretraining, posttraining A, postsleep and posttraining B test sessions.

Shortly after the completion of the posttraining A test session, the electrodes for PSG were attached (see ***PSG measurement***). Then, the subjects entered the MRI scanner. Before MRS measurement, an anatomical structure measurement, voxel placement, and shimming were conducted (see ***MRS acquisition and analysis***). The room lights were turned off, and the sleep session began.

#### Experimental design for Experiments 2 and 4

There were two test sessions: pretest and posttest sessions. The pretest session was performed before the first training session on Task A, followed by the first training session on Task A. Then, there was a 120-min interval during which the sleep session occurred (see common procedures above). After the 120-min interval, all subjects performed the second TDT training on Task B, and the posttest session was conducted.

Eighteen out of 30 subjects were tested only with Task A in the test sessions, whereas the remaining 12 subjects were tested with both Tasks A and B. The data from the latter group of subjects (n = 12) are shown in Experiment 4 (see below).

#### Experimental design for Experiment 3

After electrodes were attached for PSG measurement (see ***PSG measurement***), the subjects entered the MRI scanner. After an anatomical structure measurement, voxel placement and shimming for the MRS measurements were conducted (see ***MRS acquisition and analysis***). The room lights were turned off, and the sleep session began.

### Texture discrimination task

TDT, a standard and widely used VPL task^38^, was used in Experiments 1, 2 and 4 (Fig. 1a).

TDT was conducted in a dimly lit room. A subject’s head and chin were restrained by a chin rest. Visual stimuli were displayed on the computer screen at a viewing distance of 57 cm. TDT consists of two tasks: the orientation task and the letter task. The orientation task was the main task, whereas the letter task was designed to control subjects’ fixation^38^. Each trial began with the presentation of a fixation point at the center of the screen (1000 ms). Then, a target display was briefly presented (17 ms), followed by a blank screen of varying duration, and then by a mask stimulus (100 ms), which was composed of randomly rotated v-shaped patterns. Subjects were instructed to fixate their eyes on the center of the display throughout the stimulus presentation.

The size of the target display was 19°, which contained a 19 x 19 array of background lines. Each background line was jittered by 0.2°. The orientation of background lines was either horizontal or vertical. Each target display had 2 components: a letter (either ‘L’ or ‘T’) presented at the central location of the display and 3 diagonal lines presented in a peripheral location within a trained visual field quadrant at a 5–9° eccentricity. The target array of three lines was aligned either horizontally or vertically on the background. The location of the target array was either in the left or right upper visual field quadrants, randomly assigned to each subject, and kept consistent throughout the experiment. After the mask display, subjects used a keyboard to report whether the central letter was ‘L’ or ‘T’ (the letter task) and whether the target array was aligned ‘horizontally’ or ‘vertically’ (the orientation task). After the subject’s responses for the letter and orientation tasks, a feedback sound was delivered to indicate whether the letter task was correct (1000 Hz pure beep) or incorrect (400 Hz pure beep). No feedback was given for responses on the orientation task.

The time interval between the target onset and mask is referred to as the stimulus-to-mask onset asynchrony (SOA). The SOA was modulated across trials to control the task difficulty. The task difficulty increased with shorter SOAs.

There were 5 or 6 SOAs ranging from 316-33 ms in an experiment. In a test session, there were 15 or 20 trials for each SOA. Thus, there were a total of 75 or 120 trials in a test session. The presentation order of the SOAs was pseudorandomized to reduce the amount of learning and fatigue during the session^83^. One test session took approximately 6-10 min. In a training session, 60 trials were blocked by SOA. The order of the SOA presentation was in a descending manner. Training for Task A was conducted before sleep with one or two blocks per SOA used, with a duration of approximately 20-40 min. Training for Task B was conducted after sleep with one block per SOA, taking approximately 20 min. The orthogonal orientations (horizontal or vertical) of background lines were used for tasks A and B. Within a test or training session, there was a 10-sec break every 15 or 20 trials. Between sessions, there was a 2-min break.

Because all the subjects in Experiments 1, 2 and 4 were naive to TDT, we explained how to perform TDT in an introductory session before the first test session (pretraining test or pretest) in the experiments. During the introductory session, 3 long SOAs (800, 600, 400 ms) were used. The introductory session started with the longest SOA (800 ms) and was followed by 600-ms and 400-ms SOAs in a blocked fashion where a set of 15-20 trials was conducted for each SOA (a total of 45 or 60 trials). The introductory session was repeated until the participant performed the orientation task with at least 93% accuracy for 400-ms SOA trials.

In Experiment 1, there was a reminder session before the postsleep test session to remind subjects how to conduct the task after the 2-hr time interval. The reminder session had the same 3 sets of SOAs as the introductory session. We did not conduct a reminder session in Experiment 2 or 4 due to temporal proximity between the posttest and training session on Task B.

The threshold SOAs were obtained for each test session for each subject as follows. The percentage of correct responses for the orientation task was calculated for each SOA in a test session. A cumulative Gaussian function was fitted to obtain a psychometric curve to determine the threshold SOA that corresponds to 80% correct performance using the psignifit toolbox (ver. 2.5.6) with MATLAB (http://bootstrap-software.org/psignifit;84). All trials in which the letter task was incorrect were removed from calculation of the threshold SOA.

TDT performance changes were computed based on relative changes in the threshold SOAs (ms) between test sessions. The performance change (%) at a test session was calculated as [100 x (the threshold SOA at a previous test session – the threshold SOA at a current session) / (the threshold SOA at a previous session)].

Stimuli were generated by MATLAB software with psychotoolbox^85, 86^.

### Sleepiness measurement

The SSS rating^77, 78^ ranged from 1 (Feeling active, vital, alert, or wide awake) to 7 (No longer fighting sleep, sleep onset soon; having dream-like thoughts). Subjects were asked to choose the scale that described their state of sleepiness.

The PVT was implemented with open-source Psychology Experiment Building Language (PEBL) software^79^. In a trial of the task, after a fixation screen, a target screen was presented in which a red circle appeared in the center of the screen. The subjects were required to press the spacebar on a keyboard as quickly as possible upon detection of the circle. The time interval between the fixation and the screen with a red circle varied between 1000–4000 ms. The PVT lasted approximately for 2 min. The reaction time (sec) was log-transformed to reduce the skew of data^87^. The average reaction time was obtained as measurements of behavioral sleepiness (see **Supplementary Table 3** for sleepiness data).

### PSG measurement

#### Experiments 1 and 3: Concurrent PSG and MRS

The electrodes for PSG were attached before the introductory session for TDT, taking approximately 30 min. PSG was obtained simultaneously with structural MRI and MRS. PSG consisted of electroencephalogram (EEG), electrooculogram (EOG), electromyogram (EMG), and electrocardiogram (ECG). For 2 subjects in Experiment 1, EEGs were recorded at 18 scalp sites. For the remaining 8 subjects in Experiment 1 and all subjects in Experiment 3, EEGs were recorded at 23 scalp sites. EOGs were recorded bipolarly placed at the outer canthi of both eyes (horizontal EOG) and above and below the left and right eyes (vertical EOG). EMGs were recorded bipolarly from the mentum. ECGs were recorded from the lower shoulder blade. EEG and ECG were referenced to Fz. The ground electrode was placed at AFp4. Electrode impedances were kept at approximately or below 5 kΩ for EEG and ECG and at approximately 10 kΩ for EOG and EMG. All data were recorded by an MRI-compatible amplifier (BrainVision Recorder, Brain Products, LLC) at a sampling rate of 5000 Hz.

PSG recorded simultaneously with MRI contains two types of noise, scanner noises and ballistocardiogram artifacts. First, scanner noise was removed after the experiment using Brain Vision Analyzer 2 (Brain Products, LLC). Next, to remove ballistocardiogram artifacts, the data were low-pass filtered at 100 Hz and downsampled to 250 Hz. Then, the FMRIB plug-in for EEGLAB (The University of Oxford) was used. EEG data were rereferenced to TP9 and TP10 using EEGLAB.

#### Experiments 2 and 4: PSG only

The electrodes for PSG measurement were attached prior to the introductory session, taking approximately 45 min. PSG consisted of EEG, EOG, EMG, and ECG. EEG was recorded at 32 scalp sites, according to the 10% electrode position^88^ using active electrodes (actiCap, Brain Products, LLC) with a standard amplifier (BrainAmp Standard, Brain Products, LLC). The reference, which was Fz online, was rereferenced to the average of the left (TP9) and right (TP10) mastoids after the recording for analysis. The sampling frequency was 500 Hz. The impedance was kept below 20 kΩ. The active electrodes included a new type of integrated impedance converter, which allowed them to transmit the EEG signal with significantly lower levels of noise than traditional passive electrode systems. The data quality with active electrodes was as high as 5 kΩ using passive electrodes^75^. The passive electrodes were used for EOG, EMG, and ECG (BrainAmp ExG, Brain Products, LLC). Horizontal EOG was recorded using two electrodes placed at the outer canthi of both eyes. Vertical EOG was measured using 4 electrodes 3 cm above and below both eyes. EMG was recorded from the mentum (chin). ECG was recorded from two electrodes placed at the right clavicle and the left rib bone. The impedance was maintained at approximately 10 kΩ for the passive electrodes. The Brain Vision Recorder software (Brain Products, LLC) was used for recording. The data were filtered between 0.1 and 100 Hz. PSG was recorded in a soundproof and shielded room.

### Sleep-stage scoring and sleep parameters

Sleep stages were scored for every 30-s epoch, following the standard criteria^1, 25^ into stage wakefulness (stage W), nonrapid eye movement (NREM) stage 1 sleep (stage N1), NREM stage 2 sleep (stage N2), NREM stage 3 sleep (stage N3), and stage REM sleep (REM sleep).

Standard sleep parameters were obtained (**Supplementary Table 2**) to indicate a general sleep structure for each experiment. Sleep parameters included the sleep-onset latency (SOL, the latency to the first appearance of stage N2 from the lights off), the percentage of each sleep stage, wake time after sleep onset (WASO), sleep efficiency (SE, the total percentage spent in sleep), and the time in bed (TIB, the time interval between lights off and lights-on)^25^. Only the data from the first sleep cycle were included for the analysis.

### MRS acquisition and analysis

Subjects were scanned using a 3T Siemens Prisma scanner (Siemens) with a 64-channel head coil. It was important for subjects to sleep without discomfort and head motion during the MRI measurements. Cushions and gauze were used to stabilize subjects’ heads to reduce discomfort, and we ensured that there would be no space left between subjects’ head and the head coil to reduce head motion. A thin back cushion and knee cushion were used upon the subjects’ request. Several blankets were used to keep subjects warm and to initiate sleep during the scan.

For all MRS experiments and anatomical (T1-weighted) reference image dataset was acquired using an MPRAGE sequence (256 slices, voxel size=1×1×1 mm, 0 mm slice gap) to localize the voxel of interest (VOI) for MRS. Based on the measured anatomical structure, the VOI was manually placed on the most posterior part of the occipital lobe, covering the calcarine sulci bilaterally that corresponds to early visual areas^30^. We carefully placed the VOI in a way to include the least volume of white matter, as lipids in the white matter cause noise in spectra. Shimming for MRS was then performed using a vendor-provided automated tool (defined by the full-width-at-half maximum of the water peak; see ***Quality tests for MRS data***). The procedure for the structural measurement, VOI placement, and shimming took approximately 20-30 min.

The MEGA-PRESS sequence^89–91^ was used to measure concentrations of both GABA and Glx simultaneously from the VOI so that GABA and Glx were acquired from the same scan during the same sleep stages, according to the procedure used in the previous studies^92–97^. In addition, the concentrations of GABA and Glx were obtained from the short 2-min or 3.3-min segments (see below). It was crucial for the present study to compute the concentrations of GABA and Glx at this temporal resolution because sleep stages could shift within a few minutes.

For all the subjects in Experiment 1 and four subjects in Experiment 3, we first conducted a quick test scan of the MEGA-PRESS sequence (TR = 1.25 s, TE = 68 ms, number of average = 32, for edit-on and edit-off, see below) with double-banded pulses to simultaneously suppress the water signal^98^ and edit the γ-CH2 resonance of GABA at 3.0 ppm^30, 89, 91, 99^ to check whether the spectrum was acceptable. Second, one unsuppressed water spectrum was acquired (TR = 1.25 s, TE = 68 ms, number of average = 16) as a standard water concentration reference for single-voxel proton MRS^100–102^. Third, the 10-min water-suppressed MEGA-PRESS sequence (TR = 1.25 s, TE = 68 ms, number of average = 240 for each edit-on and edit-off, VOI = 2.2×2.2×2.2 cm^3^) was run repeatedly until the sleep session was over.

We postprocessed the raw data (twix files) from each MRS run to extract the 2-min segments for subsequent processing to obtain GABA and Glx concentrations. Since the acquisition time of the sequence was shorter than that in some previous studies^30^, we used the shorter TR and larger VOI size to increase signal-to-noise ratios.

For the remaining 6 subjects in Experiment 3, each of the acquisition times for the MEGA-PRESS scans with water suppression was 3.3 min (TR = 1.5 s, TE = 68 ms, number of average = 64, for each edit-on and edit-off, VOI = 2×2×2 cm^3^, scan time 198 s including 6 s dummy scans). None of the quick test scans or water-unsuppressed scans were performed. The spectra were obtained from the 3.3-min segment.

For all the data in Experiments 1 and 3, the final spectra were obtained by subtracting the signals from alternate scans with the selective double-banded pulse applied at 4.7 and 7.5 ppm (edit-off) and the selective double-banded pulse applied at 1.9 and 4.7 ppm (edit-on). The differential spectrum shows only the outer lines of the GABA triplet at 3.0 ppm as the creatine singlet would cancel it out. The bandwidth of the frequency selective pulses permits assessment of N-acetylaspartic acid (NAA) and glutamate resonance at 3.7 ppm.

Spectroscopic (segmented) data were processed using LCModel^103, 104^ for metabolite quantification, including Glx, GABA, and NAA. Note that Glx is a combined signal from glutamate and glutamine. The LCModel assumes that the obtained spectrum can be fitted in the frequency domain using a linear combination of basis functions. The amounts of GABA and Glx were normalized by (i.e., divided by) the amount of NAA and referred to as the concentrations of GABA and Glx, respectively^47, 93, 97, 105^. NAA was selected as a reference metabolite in our study because creatine was not obtained from the MEGA-PRESS sequence.

We assigned a sleep stage to an MRS segment in the following way. Sleep stages were scored every 30 s, while the average E/I ratio was obtained per 2-min or 3.3-min. We assigned a sleep stage to an individual MRS segment by taking the mode of sleep stages that occurred during the segment. For example, if a 2-min MRS segment included 3 epochs of NREM sleep stage 2 and one NREM sleep stage 3, NREM sleep stage 2 was assigned to the MRS segment. If the same numbers of different sleep stage epochs were included in a segment, the lighter sleep stage was assigned to the segment so that overestimation of sleep depth would not occur. If the number of EEG epochs that included noises, or if the number of arousal^25^ were equal to or more than half of all the epochs in an MRS segment, the MRS segment was removed from further analyses, as the sleep state might be unreliable due to noise or too unstable to be classified to a particular sleep stage.

The E/I ratio was calculated for each MRS segment as the Glx concentration divided by GABA concentration. Then, E/I ratios were averaged for each sleep stage, normalized by the average E/I ratio during wakefulness before sleep onset. If there was no wakefulness before sleep onset (for example, if a subject had already reached NREM sleep stage 1 when lights were turned off), wakefulness that occurred after sleep onset was used. These normalized amounts of the E/I ratio were referred to as the E/I ratio during NREM sleep [(E/I_NREMsleep_ – E/I_wake_) / E/I_wake_] × 100% and the EI ratio during REM sleep [(E/I_REMsleep_ – E/I_wake_) / E/I_wake_] × 100%, respectively.

### Quality tests for MRS data

We used a relatively short MRS segment to extract a spectrum. We conducted 4 types of quality checks, including shim values, NAA linewidth, Cramer-Rao lower bounds, and frequency drift, for the spectra described below. The results indicate that the quantification of Glx and GABA is reliable for both Experiments 1 and 3. See **Supplementary Table 5** for more details.

### Statistical analyses

The α level (Type I error rate) of 0.05 was set for all statistical analyses. The Shapiro-Wilk test was conducted for all the data to test whether the data were normally distributed. If data were not normally distributed, nonparametric tests (Mann-Whitney U test for SSS and Kruskal-Wallis test for sleep parameters) were used. Levene’s test was conducted to test for homogeneity of variance. It was confirmed that homogeneity of variance was not violated for all the data (all *p* > 0.05). The Grubbs test was conducted to detect outliers. No outlier data were included in the results.

To analyze performance improvement and MRS data, we first conducted ANOVA and performed *t*-tests as post hoc tests if necessary. Pearson’s correlation was computed to test whether there was a significant correlation between the MRS data and behavioral performances on TDT.

Statistical tests were conducted by SPSS (ver. 22, IBM Corp.) and MATLAB (R2014a, MathWorks, Inc.).

## Acknowledgments

This work was supported by NIH (R21EY028329, R01EY019466, R01EY027841, T32EY018080, and T32MH115895) and BSF2016058. Part of this research was also supported by the Center for Vision Research, Brown University.

## Author contributions

MT and YS designed the research. MT, ZW, TBD, AVB, and EW performed the experiments. MT and EW analyzed the data. All authors wrote the manuscript.

## Supplementary Materials

Supplementary Figs 1-4

Supplementary Tables 1-5 Supplementary References

