## Supplemental file for "Opponent neurochemical and functional processing in NREM and REM sleep in visual learning"

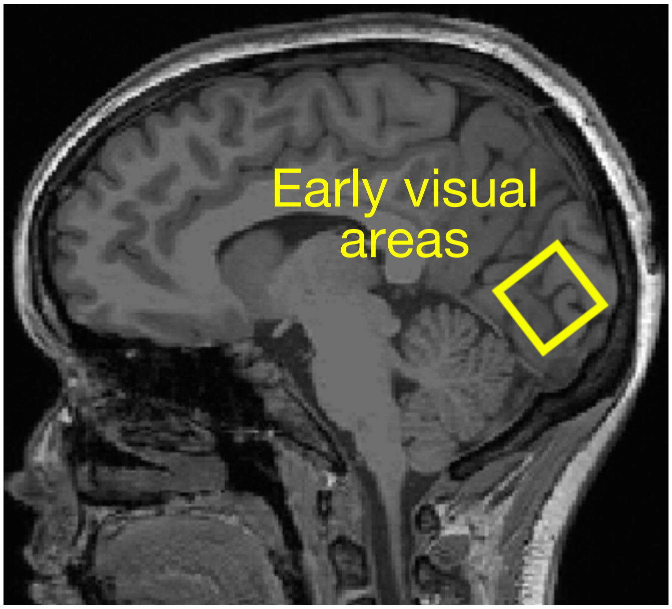


**Supplementary Fig. 1.** Example structural MRS image indicating the voxel located in early visual areas.

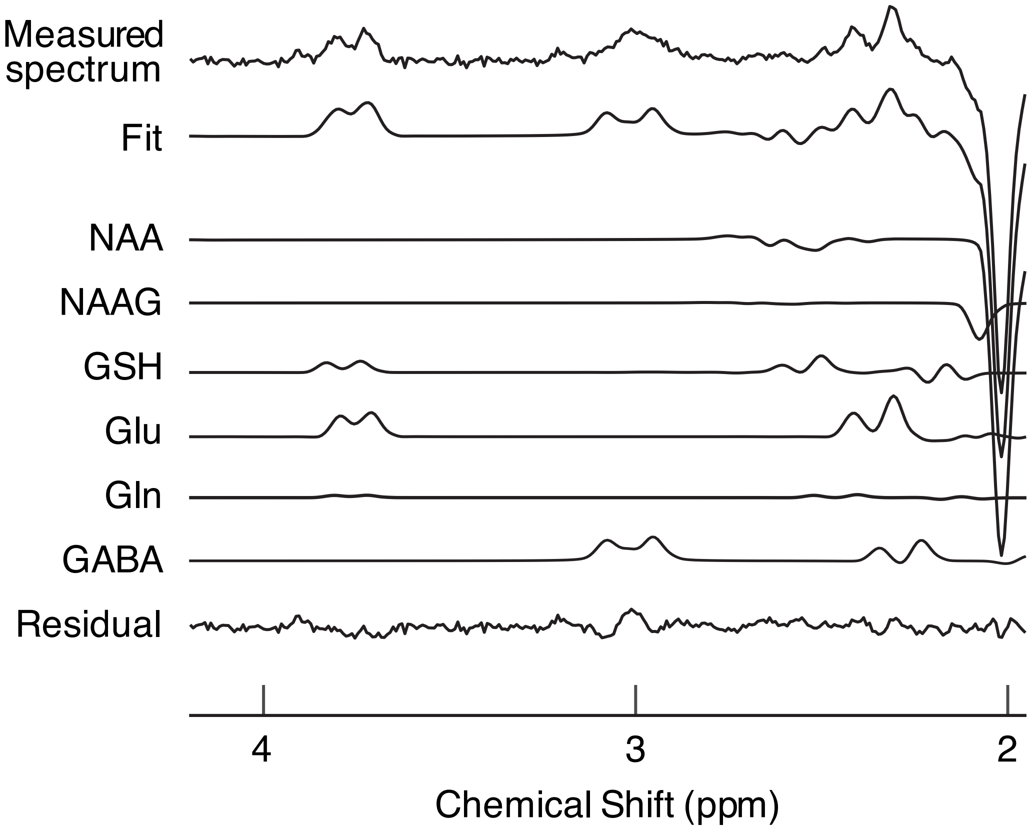


**Supplementary Fig. 2.** Example spectra from the voxel located in early visual areas. The measured spectrum is shown in the top row. “Fit” in the second row represents the spectrum fitted with the LCModel (see **MRS acquisition and analysis** in **Methods** for details). “Residual” at the bottom row represents the residual remaining after the fitting. The remaining rows show individual fits for all metabolites that can be detected by a given acquisition. Macromolecular and lipid signals were used to produce the baseline correction. NAA, NAAG, GSH, Glu, Gln, GABA represent N-acetylaspartate, N-acetylaspartylglutamate, glutathione, glutamate, glutamine, and gamma-aminobutyric acid, respectively. Glx is obtained by adding glutamine and glutamate in the LCModel.

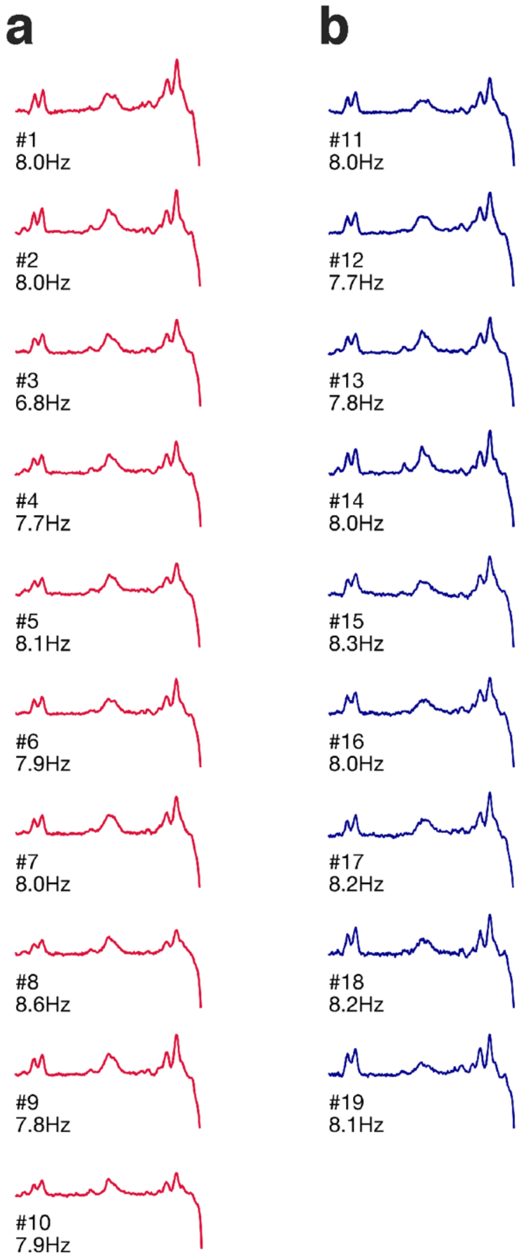


**Supplementary Fig. 3.** Mean raw spectra in Experiment 1 (**A**, n=10, red plots) and in Experiment 3 (**B**, n=9, blue plots). The value below each plot shows the mean (± SEM) full-width-at-half-maximum linewidth for NAA in Hz (Experiment 1, 7.9 ± 0.14 Hz; Experiment 3, 8.0 ± 0.07 Hz).

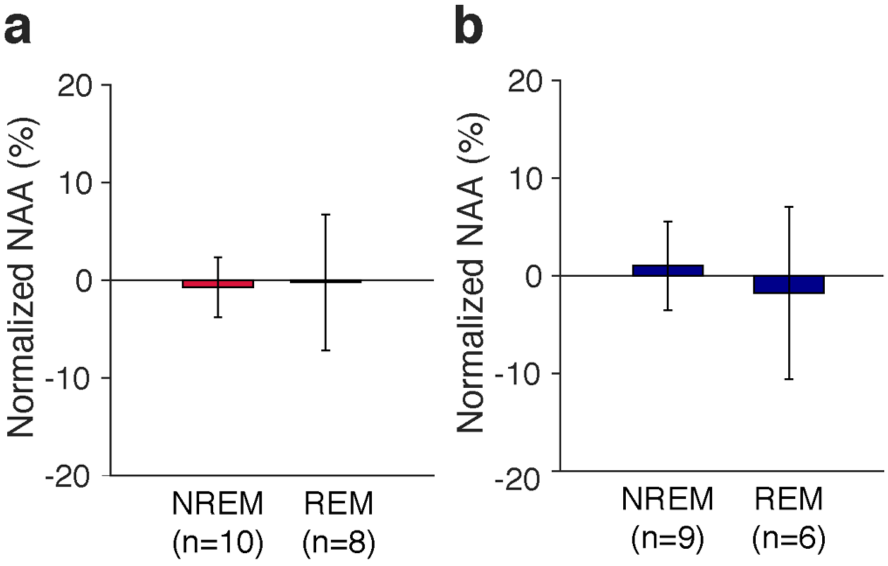


**Supplementary Fig. 4.** Results of additional analyses of MRS data. NAA concentrations during NREM sleep and REM sleep were obtained and normalized to the NAA concentrations during the awake period by $[(NAA_{sleep}-NAA_{wake})/NAA_{wake}] \times 100\%$. (**A**) Mean (± SEM) percent changes in the NAA concentrations during NREM and REM sleep for Experiment 1. (**B**) Mean (± SEM) percent changes for Experiment 3. We did not find any evidence that experiments or sleep stages significantly affected NAA concentrations as results of two-way ANOVA on NAA concentrations with the factors Experiment (Exps 1 vs. 3) and Sleep stage (NREM vs. REM sleep) showed no significant main effect of Experiment ($F$ (1, 29) = 0.001, $p$ = 0.982), Sleep stage ($F$ (1, 29) = 0.04, $p$ = 0.841) or interaction between the two factors ($F$ (1, 29) = 0.09, $p$ = 0.773). In addition, no significant change in the NAA concentration from the baseline was found for NREM sleep or REM sleep in either condition, suggesting that NAA concentrations do not significantly change between wakefulness and sleep.

**Supplementary Table 1.** Twelve subjects in Exp. 2 also participated in Exp. 4 where performances for both Tasks A and B were measured.

| Experiment | The number of subjects analyzed | The number of females | Age (mean ± SEM) | Data omission |
|---|---|---|---|---|
| 1 | 10 | 9 | 24.8 ± 0.93 | 0 |
| 2 | 30 | 18 | 22.7 ± 0.61 | 1 |
| 3 | 9 | 3 | 25.2 ± 1.21 | 1 |
| 4 | 12 | 9 | 23.2 ± 0.86 | 0 |

**Supplementary Table 2.** Sleep parameters for Experiments 1-4. We tested whether the sleep quality of each experiment was different. Since the Shapiro-Wilk test showed violation of normality for all the variables, we used Kruskal-Wallis one-way ANOVA, a nonparametric test, for each sleep parameter to test whether a sleep parameter significantly differed by experiment. We included only Experiments 1, 2, and 3, excluding Experiment 4, as subjects in Experiment 4 overlapped with those in Experiment 2. The statistical results indicated no significant difference in any of the sleep parameters across experiments. See the table for statistical results. All sleep parameters except TIB were obtained from the first sleep cycle.

| | **Exp. 1 (n=10)** | **Exp. 2 (n=30)** | **Exp. 3 (n=9)** | **Exp. 4 (n=12)** | **Kruskal-Wallis (df=2)** | |
|---|---|---|---|---|---|---|
| | | | | | **Chi square** | **P value** |
| SOL (min) | 5.0 ± 0.87 | 8.5 ± 1.00 | 6.6 ± 2.23 | 6.8 ± 1.00 | 4.47 | 0.107 |
| WASO (min) | 6.3 ± 2.64 | 5.7 ± 1.75 | 8.3 ± 4.06 | 8.3 ± 3.66 | 2.52 | 0.284 |
| Stage W (%) | 16.0 ± 3.83 | 14.2 ± 2.54 | 19.8 ± 5.89 | 16.2 ± 5.03 | 1.92 | 0.383 |
| NREM (%) | 74.6 ± 4.25 | 76.9 ± 2.12 | 75.2 ± 5.03 | 75.4 ± 3.90 | 1.88 | 0.390 |
| REM (%) | 9.4 ± 2.21 | 8.9 ± 1.46 | 5.0 ± 1.99 | 8.4 ± 2.21 | 0.06 | 0.972 |
| SE (%) | 84.0 ± 3.84 | 85.8 ± 2.54 | 80.2 ± 5.89 | 83.8 ± 5.03 | 1.92 | 0.383 |
| TIB (min) | 87.7 ± 4.14 | 87.7 ± 1.32 | 83.2 ± 6.56 | 85.0 ± 2.61 | 3.61 | 0.165 |

**SOL**, sleep-onset latency. **WASO**, wake after sleep onset. **SE**, sleep efficiency. **TIB**, the time in bed which indicates the duration of each sleep session (the time interval between lights off and lights-on). NREM sleep includes NREM sleep stages 1-3. Values are mean ± SEM.

**Supplementary Table 3.** Sleepiness data for each experiment. SSS, Stanford sleepiness scale. RT (reaction time), behavioral sleepiness measured by the reaction times (log-transformed) obtained from a psychomotor vigilance task (PVT). RTs were log-transformed. See **Sleepiness measurement** in the **Methods** for details. SSS was measured from 4 subjects, and PVT was not conducted in Exp. 3. Values are the mean ± SEM. We tested whether there was a significant difference between groups (NREM+REM vs. NREM only) for each of the experiments 1, 2, and 4. Since the Shapiro-Wilk test showed violation of normality for SSS values, we used a Mann-Whitney U test for each test session. For the RT measures, because the data were normally distributed, we performed a mixed-design ANOVA with Group as a between-subject factor and Test session as a within-subject factor. There was no significant difference between groups in SSS or RT in each experiment. See each table for statistical results.

**Experiment 1**

| Condition | | Pretraining | Posttraining A | Postsleep | Posttraining B |
|---|---|---|---|---|---|
| NREM+REM (n=8) | SSS | 1.5 ± 0.19 | 1.8 ± 0.16 | 1.1 ± 0.13 | 1.1 ± 0.13 |
| | RT | 2.55 ± 0.02 | 2.55 ± 0.01 | 2.54 ± 0.01 | 2.55 ± 0.01 |
| NREM only (n=2) | SSS | 1.5 ± 0.50 | 1.0 ± 0.00 | 1.0 ± 0.00 | 1.0 ± 0.00 |
| | RT | 2.58 ± 0.03 | 2.57 ± 0.03 | 2.52 ± 0.01 | 2.54 ± 0.02 |

The results showed no significant difference in SSS (Pretraining, U=8, p=1.000; Posttraining A, U=2, p=0.267; Postsleep, U=7, p=1.000; Posttraining B, U=7, p=1.000) or RT (Group, $F(1,8)=0.02$, $p=0.896$; Test session, $F(3,24)=3.53$, $p=0.030$; Group x Test session, $F(3,24)=0.94$, $p=0.438$) between the groups in any of the test sessions.

**Experiment 2**

| Condition | | Pre | | | Post | | |
|---|---|---|---|---|---|---|---|
| NREM+REM (n=22) | SSS | 2.0 | ± | 0.10 | 1.5 | ± | 0.11 |
| | RT | 2.51 | ± | 0.01 | 2.51 | ± | 0.01 |
| NREM only (n=8) | SSS | 1.6 | ± | 0.18 | 1.8 | ± | 0.25 |
| | RT | 2.50 | ± | 0.01 | 2.48 | ± | 0.01 |

The results showed no significant difference in SSS (Pre, U=62, z=1.55, p=0.122; Post, U=68, z=1.04, p=0.299) or RT (Group, F(1,28)=1.00, p=0.326; Test session, F(1,28)=1.76, p=0.195; Group x Test session, F(1,28)=0.38, p=0.541) between the groups in any of the test sessions.

**Experiment 3**

| Condition | | Presleep | | |
|---|---|---|---|---|
| Sleep (n=4) | SSS | 1.5 | ± | 0.29 |

| **Experiment 4** | | | | | | | |
|---|---|---|---|---|---|---|---|
| Condition | | Pre | | | Post | | |
| NREM+REM (n=9) | SSS | 2.0 | ± | 0.17 | 1.8 | ± | 0.15 |
| | RT | 2.51 | ± | 0.01 | 2.51 | ± | 0.03 |
| NREM only (n=3) | SSS | 1.7 | ± | 0.33 | 1.3 | ± | 0.33 |
| | RT | 2.53 | ± | 0.01 | 2.49 | ± | 0.01 |

The results showed no significant difference in the SSS score (Pre, U=9.5, p=0.746; Post, U=7.5, p=0.473) or RT (Group, F(1,10)=0.02, p=0.898; Test session, F(1,10)=2.08, p=0.180; Group x Test session, F(1,10)=1.40, p=0.264) between the groups in any of the test sessions.

**Supplementary Table 4.** Initial performance. Threshold SOA (ms) for TDT.

| | Condition | | Mean ± SE | Unpaired *t*-test (NREM+REM vs. NREM only) |
|---|---|---|---|---|
| Exp. 1 | NREM+REM (n=8) | | 145.0 ± 13.31 | t(8)=0.04, p=0.969 |
| | NREM only (n=2) | | 143.9 ± 5.91 | |
| Exp. 2 | NREM+REM (n=22) | | 138.8 ± 9.83 | t(28)=0.95, p=0.349 |
| | NREM only (n=8) | | 160.6 ± 27.16 | |
| Exp. 4 | NREM+REM (n=9) | Task A | 145.3 ± 19.49 | Task A: t(10)=1.46, p=0.176 |
| | | Task B | 183.7 ± 22.82 | |
| | NREM only (n=3) | Task A | 211.6 ± 56.73 | Task B: t(10)=0.06, p=0.954 |
| | | Task B | 186.5 ± 44.71 | |

**Supplementary Table 5.** MRS data quality.

| | Shim value (Hz) | NAA linewidth (Hz) | Frequency drift (Hz) | %SD for Glx | %SD for GABA |
|---|---|---|---|---|---|
| Experiment 1 | 14.0 ± 0.19 | 7.9 ± 0.14 | 1.1 ± 0.20 | 7.6 ± 0.22 | 9.1 ± 0.38 |
| Experiment 3 | 14.3 ± 0.28 | 8.0 ± 0.07 | 1.1 ± 0.13 | 6.3 ± 0.37 | 8.8 ± 0.41 |

**Note:**
***Shim values.*** The shim values represent the homogeneity of the magnetic field, which was obtained once for each MRS run. A shim value of less than 30 Hz is regarded as desirable[1]. In the present study, the mean shim value was much lower than 30 Hz and comparable to a previously reported value[2].
***NAA linewidth.*** The linewidth for NAA, which determines the resolution available to discern spectral features (therefore, the lower the better), was noted for each MRS run for each subject. We confirmed that the NAA linewidth was all below 10 Hz (see **Fig. S3** for the NAA linewidth, and see **Fig. S2** for example spectra). The mean NAA linewidth for each experiment was comparable to the values reported in previous studies[2,3].
***Frequency drift.*** We measured the frequency drift[2,4], in which larger values suggest head motions. The mean frequency drift values were in an acceptable range, similar to the previous studies[2,3]. There were no frequency drift data from the 5 subjects in Experiment 3. Thus, the frequency drift in Experiment 3 was calculated from 4 subjects.
***Cramer-Rao lower bounds.*** The Cramer-Rao lower bounds (or %SD) were used as a measure of fitting errors (therefore, the lower the better), and a commonly accepted Cramer-Rao lower bound criterion of 20% was chosen to reject low-quality signal[5]. The mean %SD values for Glx and GABA were similar to the reported values in previous studies[3].
***Comparison of the MRS quality between Experiments 1 and 3.*** To test whether MRS data quality was significantly different between Experiments 1 and 3, MANOVA with a factor of Experiment was conducted with shim values, NAA linewidth, %SD for Glx and %SD for GABA as the dependent variables. Box's test of equality of covariance matrices showed that Box's M value was not significant ($F$ (10, 1339.5) = 1.72, $p$ = 0.071), indicating the homogeneity of variance-covariance. MANOVA showed no significant main effect of Experiment (*Pillai's Trace = 0.37*, $F$ (4, 14) = 1.97, $p$ = 0.155). Note that the frequency drift was not included in MANOVA due to lack of data. For the frequency drift, we conducted unpaired $t$-test between Experiments 1 and 3. There was no significant difference between the experiments in the frequency drift ($t$ (12) = 0.08, $p$ = 0.937). Overall, the MRS data quality is reasonable and comparable to previous studies, with no significant difference between Experiments 1 and 3.
